# Astrocyte gap junctions and Kir channels contribute to K^+^ buffering and regulate neuronal excitability

**DOI:** 10.1101/2025.02.10.637549

**Authors:** Danica Bojovic, Andre Dagostin, Steve J. Sullivan, Henrique von Gersdorff, Anusha Mishra

## Abstract

Astrocytes are connected in a functional syncytium via gap junctions, which is thought to contribute to maintenance of extracellular K^+^ homeostasis. The prevailing hypothesis is that K^+^ released during neuronal firing is taken up by astrocytes via K_ir_ channels and then distributed among neighboring astrocytes via gap junctions. Previous reports examining the role of K_ir_ channels and gap junctions have shown both hyperexcitability and depression when each mechanism is blocked. Here, we tested the effect of blocking K_ir_ channels and gap junctions, both independently and simultaneously, on field activity of cortical slices in response to a 3 s, 20 Hz stimulation train. Independently blocking either K_ir_ channels or gap junctions increased the amplitude of the first fEPSC (field excitatory post-synaptic current) in response to a stimulation train, followed by suppression of fEPSCs during sustained stimulation. Surprisingly, blocking both gap junctions and K_ir_ channels enhanced the suppression of neuronal activity, resulting in a ∼75% decrease in fiber volley (pre-synaptic action potentials) amplitude in the first response, followed by a fast and strong suppression of sustained fEPSCs. Our results demonstrate that blocking K_ir_ channels and gap junctions can increase the excitability of neurons when firing is sparse, but suppression results when the firing frequency is increased to cortical physiological ranges. This suggest that K^+^ buffering via K_ir_ and gap junctions, likely mediated by astrocytes, together play a critical role in maintaining neuronal excitability, particularly during sustained activity.

## INTRODUCTION

Neuronal excitability relies on the maintenance of ionic gradients across cell membranes. In this context, potassium ion (K^+^) gradient is characterized by high intracellular and low extracellular concentrations. Although K^+^ homeostasis is necessary for optimal neuronal firing, how extracellular K^+^ concentration ([K^+^]_e_) is regulated is not completely understood. Presumably, [K^+^]_e_ is largely regulated by astrocytes, which express high levels of inwardly rectifying K^+^ 4.1 channels (K_ir_ 4.1) that take up the K^+^ released from active neurons^1^. Astrocytes also form a syncytium connected via gap junctions (GJs), composed of connexin 30 and 43 (Cx30 and Cx43)^2^. The prevailing hypothesis is that astrocytes take up K^+^ via K_ir_ 4.1 channels and then spread it throughout the syncytium via GJs to maintain [K^+^]_e_ homeostasis during bouts of neuronal activity^1,3^.

Changes in [K^+^]_e_ are observed during aging and neurodegeneration^4^, underscoring the importance of understanding the effect of K^+^ homeostasis on neuronal health. Studies have reported that an increase in [K^+^]_e_ leads to either depression^5,6^ or hyperexcitation^7,8^ of neuronal firing, but the reason for these disparate reports are unclear. As K_ir_4.1 channels and GJs are thought to be the key players in buffering [K^+^]_e_, many studies have investigated their roles in K^+^-dependent changes of neuronal excitability. Yet, K_ir_ channels and GJs are often studied separately, resulting in incomplete conclusions about the specific contribution of each mechanism to K^+^ buffering and their impact on neuronal excitability^1,2,9^. Furthermore, although studies using genetic manipulations to remove K_ir_4.1 channels and GJs have provided evidence that they both contribute to this process, these methods can also induce widespread compensatory or adaptive transcriptional changes^1,2^, making it difficult to interpret the findings.

We evaluated how K_ir_ and GJs affect neuronal excitability in cortical slices using acutely applied inhibitors to assess their respective contributions. In response to a moderate 3 s, 20 Hz frequency neuronal stimulation, blocking either K_ir_ and GJs independently resulted in similar initial post- synaptic hyperexcitability followed by a faster reduction in the amplitude of neuronal fiber volley and post-synaptic responses. The prevailing hypothesis would suggest that blocking K_ir_ channels first should block the initial K^+^ influx into astrocytes and subsequent GJ block would not lead to further disruption of neuronal activity. However, surprisingly, we have found that the combined block of K_ir_ and GJs resulted in a stronger reduction of both fiber volley and post-synaptic activity. Together, our data indicate that K_ir_ channels and GJs, likely in astrocytes, act in concert to orchestrate K^+^ buffering and preserve sustained neuronal firing.

## MATERIALS AND METHODS

### 2.1. Animals

All experimental procedures were performed in accordance with the Institutional Animal Care and Use Committee (IACUC) at Oregon Health & Science University. C57BL/6J mice of both sexes at 4-8 weeks of age were used.

### 2.2. Preparation of acute cortical slices

Cortical slices were prepared as described previously^10^. Mice were quickly euthanized under anesthesia (4% isoflurane) and the brain removed. Cortical slices (300 μm) were cut with a Compresstome VF-300-0Z (Precisionary Instruments Inc., USA) in oxygenated (95% O_2_/5% CO_2_) ice-cold slicing solution containing 93 mM N-methyl-D-glucamine (NMDG)-Cl, 2.5 mM KCl, 0.5 mM CaCl_2_, 10 mM MgCl_2_, 1.2 mM NaH_2_PO_4_, 30 mM NaHCO_3_, 20 mM HEPES, 25 mM D-glucose, 1 mM kynurenic acid, 5 mM Na ascorbate, 3 mM Na pyruvate (NMGD replaces Na in the slicing solution to provide charge and osmotic pressure but prevent excitotoxicity). Brain slices were placed in a warm recovery bath with the same slicing solution at 37° C for 20 min, then transferred to an oxygenated (95% O_2_/5% CO_2_) storage solution containing 92 mM NaCl, 2.5 mM KCl, 2 mM CaCl_2_, 1 mM MgCl_2_, 1.2 mM NaH_2_PO_4_, 30 mM NaHCO_3_, 20 mM HEPES, 25 mM D-glucose, 1 mM kynurenic acid, 5 mM Na ascorbate, 3 mM Na pyruvate at room temperature and allowed to equilibrate for at least 30 min before use. Experiments were performed in 1-3 slices from each animal, up to 6 hours after slicing.

### 2.3. Electrophysiology

Slices were transferred to a recording chamber and continuously perfused (3-4 ml/min) with oxygenated (20% O_2_, 5% CO_2_, 75% N_2_) artificial cerebrospinal fluid (aCSF) composed of 124 mM NaCl, 3.5 mM KCl, 2 mM CaCl_2_, 1 mM MgCl_2_, 1 mM NaH_2_PO_4_, 26 mM NaHCO_3_, 10 mM glucose, and 1 mM Na ascorbate. Slices were visualized under an upright Olympus BX51WI microscope (Japan) under DIC optics. Borosilicate glass pipettes (World Precision Instruments, USA) were pulled into glass electrodes with a Micropipette Puller (Sutter Instruments model P-87, USA). The recording electrode resistance was ∼3-5 MΩ, while the stimulating electrode had a large diameter (15 to 25 μm) tip with low resistance. Both electrodes were filled with aCSF and inserted into the region of interest, with the stimulating electrode in layers I-II and recording electrode in layers IV-V of the cortex. Field currents were recorded in voltage-clamp mode (Axon Instruments, Molecular Devices, USA). All experiments were conducted at 33-36° C. Single stimulation pulses were delivered to find the field response before starting the experiment. A 30 min stabilization period was allowed, following which a 3 s, 20 Hz stimulation train was delivered to obtain baseline (pre-treatment) for each slice, and again after 1 hour bath application of drugs of interest (post-treatment). Three traces were collected for baseline and drug conditions, with 2 min intervals between recordings, and averaged to represent each condition.

### 2.4. Pharmacology

AMPA receptors were blocked by 10 μM cyanquixaline (CNQX; Tocris Bioscience, UK); voltage-gated Na^+^ channels with 1 μM tetrodotoxin (TTX; Tocris Bioscience, UK); gap junctions with 100 μM meclofenamic acid (MFA; Sigma Aldrich, USA) and 100 μM carbenoxolone (CBX; Tocris Bioscience, UK); and K_ir_ channels with 200 μM BaCl_2_ (Sigma Aldrich, USA).

### 2.5. Data analysis

Field recordings were analyzed in Igor Pro with Neuromatic extension (Wave Metrics, USA). Statistical analysis was performed in Prism (Graph Pad, USA) and Excel (Microsoft, USA). Data are represented as mean ± SEM. The significance level was set at p < 0.05.

All traces were first smoothed with binomial function. Peak amplitude was calculated as the difference between peak of the fiber volley (pre-synaptic action potentials) or fEPSC (field post- synaptic current) and the average baseline value before stimulation. To calculate the change in amplitude after each treatment, the first peak amplitude after 1 hour was normalized to the first pre-treatment amplitude. To calculate the amplitude change over the 3 s, 20 Hz stimulation, each of the 60 traces were normalized to the first trace. Time to 50% amplitude was calculated by finding the time point when amplitude decreased below 50% of the maximal amplitude change over the 3 s stimulation. Wilcoxon test was used to compare paired pre-treatment to post-treatment values. Mann-Whitney test was used to compare unpaired samples between two groups. Kruskal-Wallis test was used for comparison of unpaired samples between multiple groups.

## RESULTS

### Experimental setup for assessing neuronal field activity

We measured evoked field activity in cortical layers IV/V in response to 3 s, 20 Hz stimulation. The recording consists of the stimulation artifact, fiber volley (pre-synaptic action potentials), and field excitatory post-synaptic currents (fEPSC; Fig. 1 A). Blocking AMPA receptors with 10 μM cyanquixaline (CNQX; Fig. 1 A) abolished the second peak, confirming it as fEPSC, while blocking voltage-gated Na^+^ channels with 1 μM tetrodotoxin (TTX; Fig. 1 A) abolished all response except the stimulation artifact, demonstrating that the first peak represents the pre-synaptic fiber volley. Field recordings were allowed to stabilize for 30 min, a period during which the signal strength steadily increased (Supp. Fig. 1). After stabilization, the 3 s, 20 Hz stimulation (Fig. 1 B) was delivered 3 times in 2 min intervals to record and quantify the neuronal activity and repeated after 1 hour treatment under different conditions (Figure 1C).

**Figure 1.**
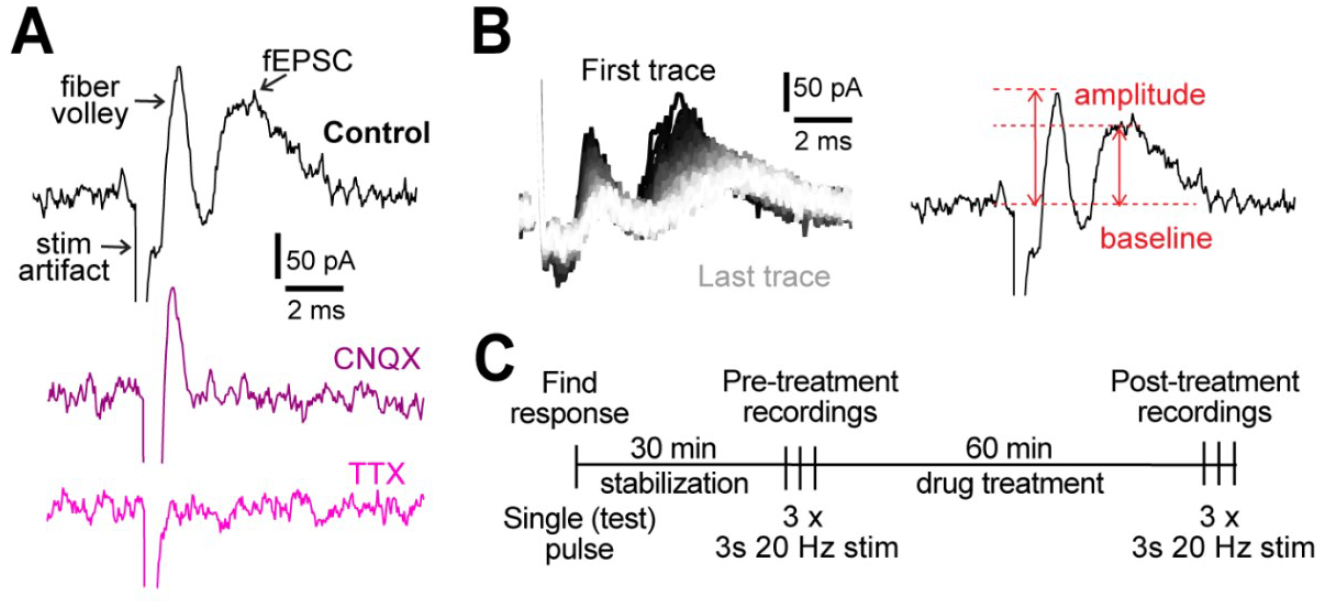
Experimental design. **A**. Representative examples of field recordings showing that the post- synaptic currents are blocked by 10 μM CNQX (purple) and fiber volley by 1 μM TTX (pink). **B**. A representative example of all 60 traces from a 3 s, 20 Hz stimulation (left). The order of traces is represented with a gradient, first (black) to last (white). Peak amplitude was measured as the difference between the highest point in fiber volley and fEPSC compared to the baseline (right). **C**. Experimental protocol used for all experiments. **D**. Table of Drugs used in this study.

Under control conditions (aCSF), we measured an increase in neuronal post-synaptic response to the 3 s, 20 Hz stimulation after 1 hour compared to the pre-treatment. The peak amplitude of the first fiber volley stayed similar: it was 93.6 ± 55.4 pA at the start and 111.8 ± 89.7 pA after 1 hour (p=0.07; Fig. 2 Ai, B). However, the peak amplitude of first fEPSC almost doubled in size from 132.1 ± 89.1 pA at the start to 234.3 ± 174.4 pA after 1 hour (p<0.0001; Fig. 2 Ai, C). While the 30 min stabilization period (Supp. Fig. 1) helped slow down the increase in fEPSC, they continued to increase in size throughout the 1 hour recording period.

**Figure 2.**
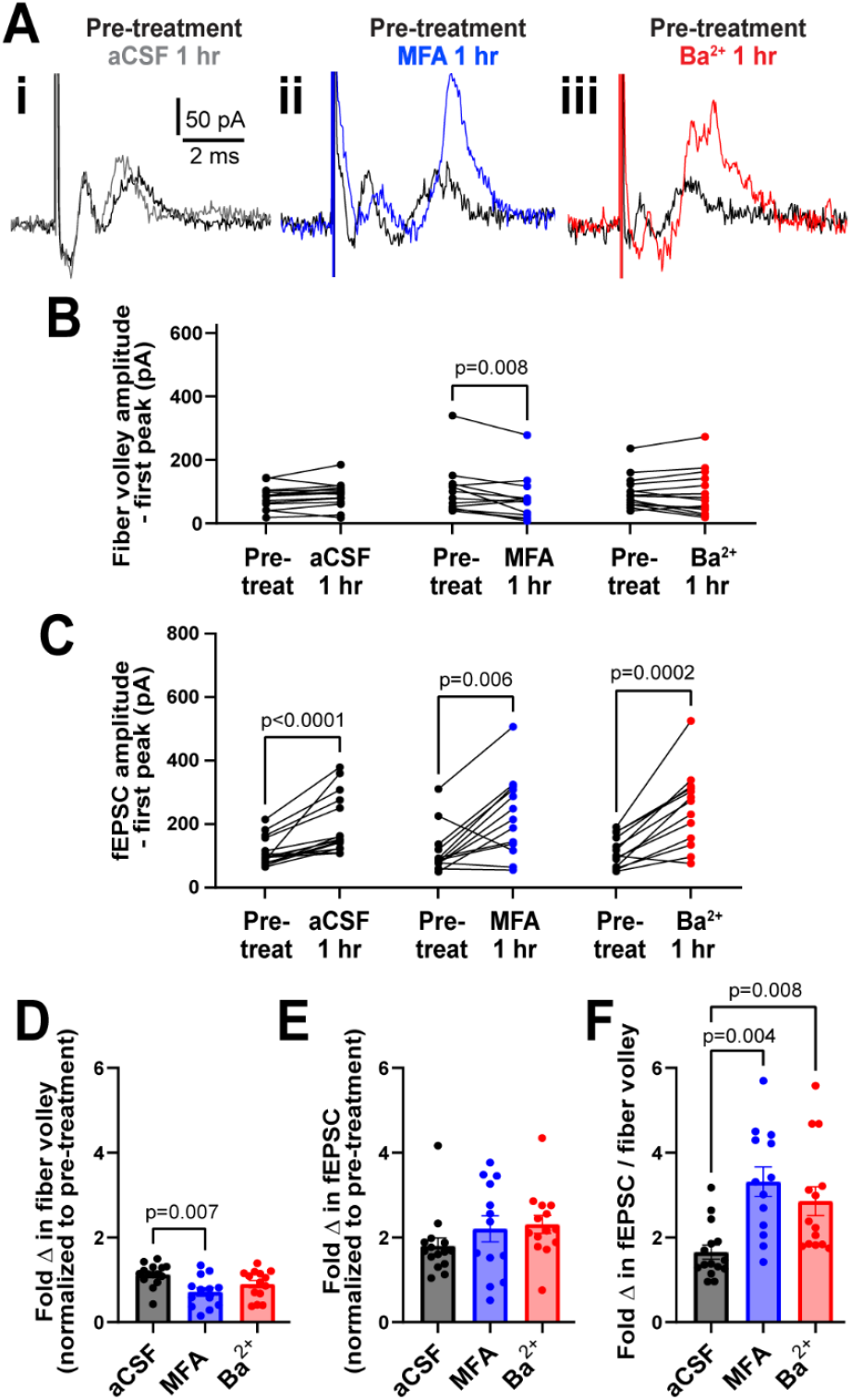
GJs and Kir channels regulate neuronal field responses. **A**. Representative traces of first recordings from slices treated with aCSF (i), the GJ blocker MFA (ii), and K_ir_ channel blocker Ba^2+^ (iii). **B**. Amplitude of the first fiber volley did not change after 1 hour in aCSF (n=15) or Ba^2+^ (n=14), but decreased in MFA (n=13, Wilcoxon test). **C**. Amplitude of the first fEPSC increased after 1 hour in all three conditions (Wilcoxon test). **D**. The fold change in the amplitude of first fiber volley f showing a relative decrease after 1 hour incubation in MFA, but not Ba^2+^, compared to aCSF. **E**. The fold change in the amplitude of the first fEPSC after 1 hour in MFA and Ba^2+^ was similar to aCSF. **F**. The fEPSC:fiber volley ratio showed a significant increase in both MFA and Ba^2+^ (D-F, Kruskal-Wallis test). Data shown as mean ± sem.

### Gap junctions and K_ir_ channels independently regulate neuronal activity

The 3s, 20 Hz stimulation was repeated before and after 1 hour treatment with meclofenamic acid (MFA; gap junction blocker) and barium (Ba^2+^; K_ir_ channel blocker) applied in the bath. To examine the effect of GJs on neuronal activity, we chose MFA, a GJ blocker shown to have higher potency than other commonly used blockers^11^. In particular, MFA blocks GJ coupling between astrocytes after 15-25 min of treatment, although not completely, resulting in increased input resistance and decreased capacitance of astrocytes^12^. In our experiments, MFA (100 μM) treatment for 1 hour reduced the peak amplitude of the first fiber volley in the stimulation train by 24.5%, from 101.5 ± 80.7 pA to 76.6 ± 72.3 pA (p=0.008, Fig. 2 Aii, B), but increased the peak amplitude of the first fEPSC by 95.5%, from 114.9 ± 73.8 pA to 224.6 ± 126.2 pA (p=0.006; Fig. 2 Aii, C). Treatment with the vehicle dimethyl sulfoxide (0.1%) alone (used for MFA treatment) did not alter neuronal excitability compared to aCSF controls (data not shown; n=9).

Next, we used the K_ir_ channel blocker barium (Ba^2+^, 200 μM) and found that it had no effect on the peak fiber volley amplitude, which was 98.6 ± 52.4 pA before treatment and 95.9 ± 72.0 pA after 1 hour (p=0.7; Fig. 2 Aiii, B), but increased the peak amplitude of the first fEPSC by 123.3%, from 112.6 ± 47.4 pA to 251.4 ± 116.1 pA (p=0.0002; Fig. 2 Aiii, C).

To compare the change in response between treatment conditions, we normalized the response size in the first trace of the stimulation train after 1 hour treatment to the pre-treatment baseline for each condition. The fold change in peak fiber volley amplitude after 1 hour of MFA treatment was smaller compared to control aCSF condition (aCSF: 1.1 ± 0.2; MFA: 0.7 ± 0.4, p=0.007; Fig. 2 D), while that after Ba^2+^ treatment was somewhat lower but not statistically significant (Ba^2+^: 0.9 ± 0.3, p=0.2, Fig. 2 D). Surprisingly, neither MFA nor Ba^2+^ affected the fold change in peak fEPSC amplitude (aCSF: 1.8 ± 0.7; MFA: 2.2 ± 1.1, p=0.4; Ba^2+^: 2.3 ± 0.8, p=0.06; Fig. 2 E).

The decrease in fiber volley despite the fEPSCs remaining similar in size suggested that the size of the post-synaptic response per pre-synaptic axon stimulated was different between conditions. Thus, we calculated the fEPSC to fiber volley ratio for each experiment. We found that the fEPSC:fiber volley ratio almost doubled in both MFA and Ba^2+^ compared to control (aCSF ratio = 1.7 ± 0.6 pA; MFA = 3.3 ± 1.3 pA, p=0.0004; Ba^2+^ = 2.9 ± 1.3 pA, p=0.008; Fig. 2 F). These data show that both K_ir_ and GJs regulate neuronal excitability in a similar manner.

There was considerable response variability in all conditions and the fEPSCs increased in size over time even under the control condition (Fig. 2 C). This variability could not be explained by age (Supp. Fig. 3 A) or sex (Supp. Fig. 3 B) of the mouse. These observations underscore the need to perform cortical field recording experiments in a paired manner in the same slices within a consistent time window to obtain meaningful data regarding drug-effects on neuronal excitability.

We also examined a second inhibitor of GJs, carbenoxolone (CBX), which is more commonly used but is perhaps less potent^11^. Slices treated with CBX (100 μM) showed a surprisingly mild effect; CBX increased peak fiber volley amplitude by 32.8 %, from 86.8 ± 34.7 pA pre-treatment to 115.3 ± 55.7 pA after 1 hour in CBX (p=0.008; Supp. Fig. 3 A) but had no statistical effect on fEPSC amplitude (pre-treatment: 106.5 ± 53.6 pA; post-treatment = 133.8 ± 68.1 pA; p=0.08; Supp. Fig. 3 B). Compared to aCSF control after 1 hour, the fold change in fiber volley amplitude (Supp. Fig. 3 C), fEPSC amplitude (Supp. Fig. 3 D), and the fEPSC:fiber volley ratio (Supp. Fig. 3 E) were all unchanged.

### Gap junctions and K_ir_ channels independently contribute to sustained neuronal activity

When examining the response to the 3s, 20 Hz stimulation train, we observed that the amplitude of the fiber volley and the fEPSCs both steadily decreased over 3 s of stimulation (Fig. 3 A). To quantify this decrease of neuronal activity, we plotted the amplitude of each fiber volley and fEPSC over the 3 s period and quantified the time it takes for amplitude to decrease below 50% of the maximum change measured at the end of the stimulation train. Time to 50% amplitude of the fiber volley decreased slightly after 1 hour but was statistically similar to the pre-treatment in all conditions (aCSF: from 1.48 ± 0.6 s to 1.0 ± 0.8 s, p=0.9; MFA: from 1.3 ± 0.3 s to 1.0 ± 1.0 s, p=0.2; Ba^2+^: from 1.4 ± 0.3 s to 1.0 ± 0.8 s, p=0.06; Fig. 3 A-D). Notably, time to 50% amplitude of fEPSC was shorter after 1 hour compared to the pre-treatment in all conditions (Fig. 3 A). In aCSF, the time to 50% amplitude of fEPSCs decreased from 0.7 ± 0.5 s to 0.4 ± 0.2 s (p=0.08), but decreased drastically in MFA, from 1.3 ± 0.8 s to 0.3 ± 0.2 s (p=0.02), and in Ba^2+^, from 1.2 ± 0.7 s to 0.2 ± 0.3 s (p=0.0002; Fig. 3 E-G).

**Figure 3.**
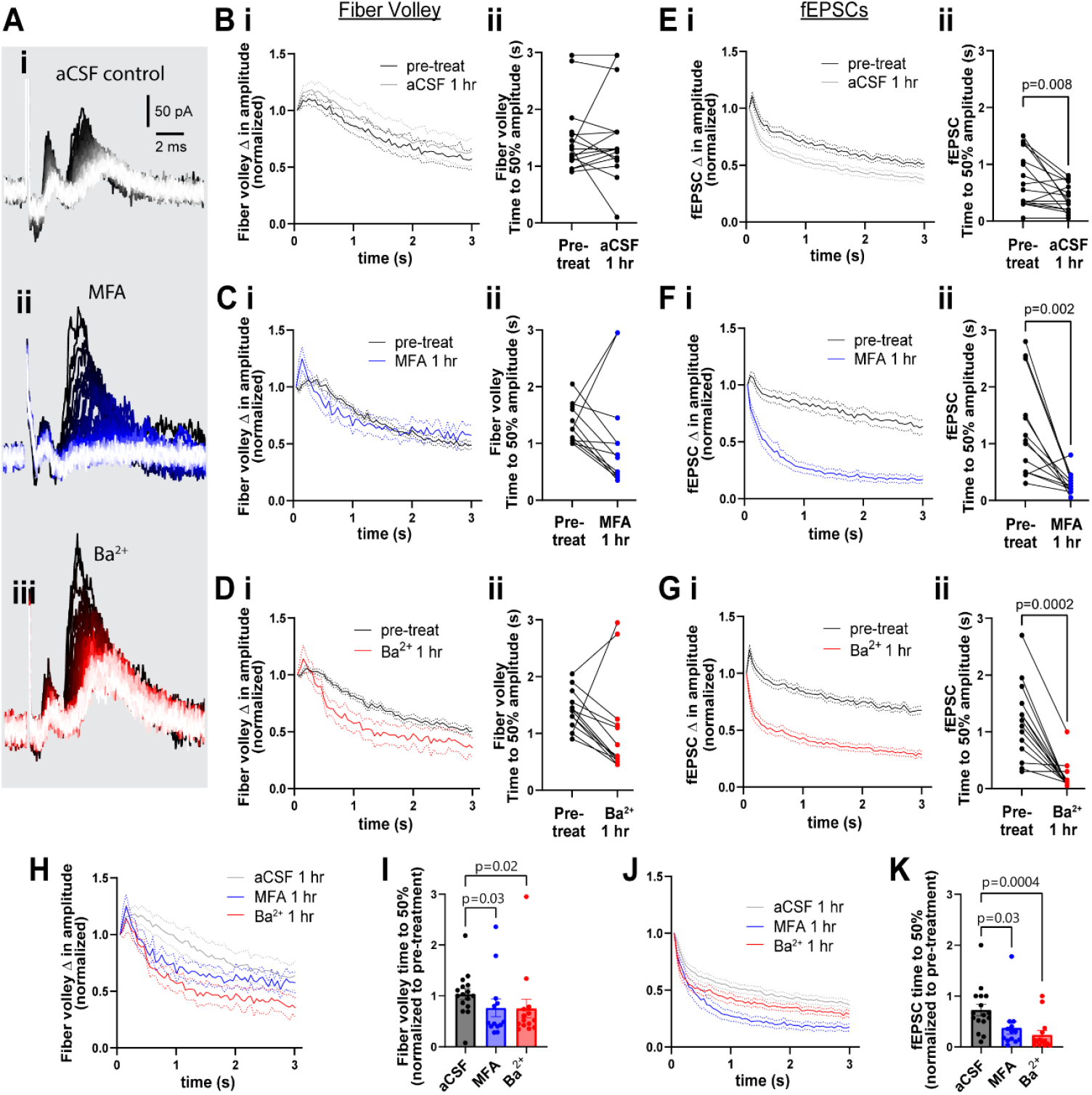
GJs and Kir channels contribute to maintenance of sustained neuronal activity. **A**. Representative traces of neuronal response to 3 s, 20 Hz stimulation in aCSF control (i), and in presence of MFA (ii), and Ba^2+^ (iii). **B-D**. Change in fiber volley amplitude over the 20 Hz stimulation (i) and time to 50% of maximum decrease (ii) showing no difference between pre-treatment and after 1 hour in aCSF (**B**; n=16), MFA (**C**; n=13), or Ba^2+^ (**D**; n=14, Wilcoxon test). **E-G**. Change in fEPSC amplitude over the 20 Hz stimulation (i) and time to 50% of maximum decrease (ii) shows a strong decrease after 1 hour in aCSF (**E**), MFA (**F**), and Ba^2+^ (**G**; n=14, Wilcoxon test) compared to pre-treatment. **H**. The decrease in fiber volley over 20 Hz stimulation after 1 hour in aCSF, MFA, and Ba^2+^ (data pooled from **B, C** and **D**, overlaid for comparison. **I**. Fold change in fiber volley time to 50% showing the relative decrease after 1 hour in MFA and Ba^2+^ (Kruskal-Wallis test). **J**. The decrease in fEPSC over 20Hz stimulation after 1 hour in control, MFA, and Ba^2+^ (data pooled from **E, F** and **G**, overlaid for comparison). **K**. Fold change in fEPSC time to 50% showing the relative decrease after 1 hour in MFA and Ba^2+^ (Kruskal-Wallis test). Data shown as mean ± sem.

Between group comparison showed that both fiber volley and fEPSC amplitudes decreased faster in presence of MFA or Ba^2+^ compared to aCSF. Fiber volley decreased 20% faster in both MFA and Ba^2+^-treated slices (fold change in time to 50% amplitude in aCSF: 1.0 ± 0.4; MFA: 0.8 ± 0.6, p=0.03, Ba^2+^: 0.8 ± 0.7, p=0.02, Fig. 3 H-I). The decrease in fEPSC was more pronounced: 42.9% faster in MFA and 71.4% faster in Ba^2+^-treated slices (aCSF: 0.7 ± 0.5; MFA: 0.4 ± 0.4, p=0.03, Ba^2+^: 0.2 ± 0.3, p=0.0004; Fig. 3 J-K).

By contrast, CBX had a weak effect on the rate of decrease of fiber volley, with time to 50% decreasing from 1.8 ± 0.7 s in pre-treatment to 1.0 ± 0.3 s after 1 hour in CBX (p=0.02; Supp. Fig. 3 F-G), and had no statistical effect on the time to 50% of fEPSC (pre-treatment: 1.4 ± 1.0 s; post-treatment: 0.8 ± 0.6 s; p=0.3; Supp. Fig. 3 H-I).

### Gap junctions and K_ir_ channels coordinate together to regulate neuronal activity

The similar effect of blocking GJs and K_ir_ channels independently suggests that they may work together to regulate neuronal activity. To examine their combined effects, we incubated slices in a cocktail of both inhibitors (100 μM MFA and 200 μM Ba^2+^) together for 1 hour and examined neuronal responses. MFA + Ba^2+^ applied together massively reduced the peak of the first fiber volley amplitude by 70%, from 73.6 ± 36.8 pA to 22.3 ± 30.4 pA (p<0.0001, Fig. 4 A, B). While MFA + Ba^2+^ had a variable effect on the peak amplitude of the first fEPSC, which was increased in some and decreased in others, the net effect was not statistically different (118.3 ± 63.1 pA pre-treatment vs. 93.5 ± 78.3 pA, p=0.2; Fig. 4 A, C). Between-group comparison showed that, compared to control, MFA + Ba^2+^ condition exhibited a much smaller fold change in the peak fiber volley amplitude (control: 1.2 ± 0.3, MFA + Ba^2+^: 0.3 ± 0.3, p<0.0001; Fig. 4 D) as well as a decrease in the fEPSC amplitude (control: 1.8 ± 0.7, MFA + Ba^2+^: 0.8 ± 0.5, p<0.0001, Fig. 4 E). The fEPSC:fiber volley ratio after MFA + Ba^2+^ (0.5 ± 0.5) was not different compared to controls (0.6 ± 0.1, p=0.07; Fig. 4 F). We suggest this effect can be attributed mainly to the large decrease in fiber volley and indicates that the fEPSC decline was likely due to fiber volley decline.

**Figure 4.**
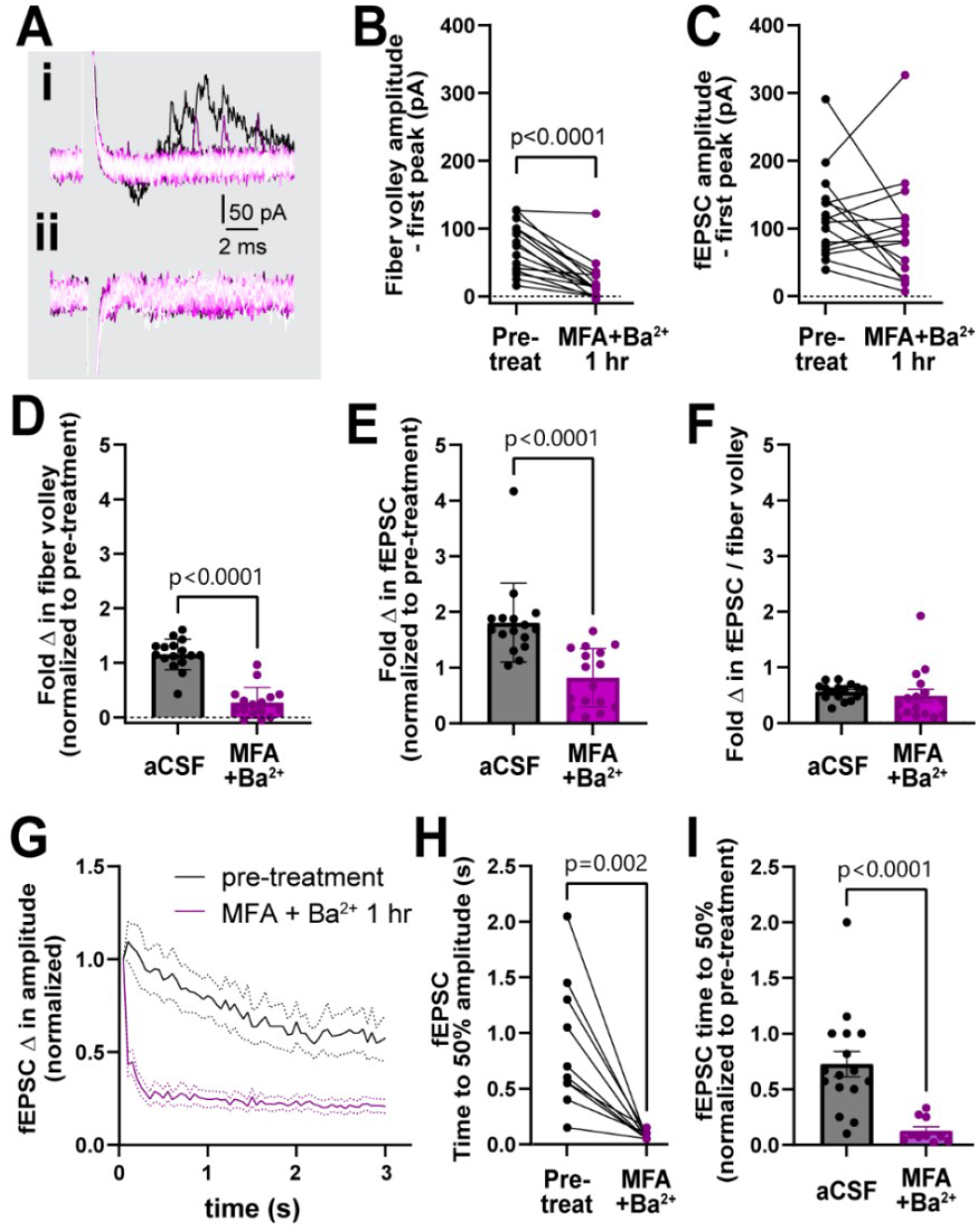
Simultaneous treatment with MFA and Ba^2+^ strongly disrupted neuronal activity. **A**. Representative traces (i, ii) of field recording during 20 Hz stimulation in presence of MFA + Ba^2+^. **B**. The amplitude of the first fiber volley decreased after 1 hour in MFA + Ba^2+^ compared to the pre- treatment (n=15, Wilcoxon test). **C**. The amplitude of the first fEPSC was not significantly different (n=16, p=0.2, Wilcoxon test). **D-E**. Fold change in fiber volley (D) and fEPSCs (E) amplitudes were smaller in MFA + Ba^2+^ compared to aCSF controls (aCSF, n=14; MFA + Ba^2+^, n=16) **F**. Fold change in fEPSC:fiber volley ratio showed no change (control n=14, Kruskal-Wallis test). **G**. Average decrease in fEPSC amplitude during the 20 Hz stimulation before and 1 hour after MFA + Ba^2+^ application. **H**. Time to 50% fEPSC amplitude was smaller in presence of MFA + Ba^2+^ compared to the pre- treatment (p<0.0001, Wilcoxon test). **I**. Time to 50% fEPSC amplitude was also smaller compared to aCSF control (Mann-Whitney test). aCSF control data are the same as in Fig. 2 D. Data shown as mean ± sem.

The fiber volley peak amplitude was so drastically low in MFA + Ba^2+^ even in the first trace that time to 50% amplitude over the course of the 3s, 20 Hz stimulation could not be accurately calculated (Fig. 4 A). However, fEPSC amplitude often spiked in the first recording followed by a sharp decrease in amplitude (Fig 4. Ai), which made it more feasible to calculate the time to 50% amplitude. The fEPSC amplitude decreased strongly, showing a decrease in time to 50% amplitude from 0.9 ± 0.6 s before treatment to 0.07 ± 0.03 s (p=0.002; Fig. 4 G-H) after 1 hour in MFA + Ba^2+^. This strong suppression of neuronal activity prompted us to record the response to stimulation train at in-between timepoints, 20 min and 40 min (Fig. 5 A). We found that time to 50% of fEPSC amplitude decreased progressively faster over the 1 hour period (pre-treatment: 0.9 ± 0.6 s; 20 min post-treatment: 0.9 ± 1.2 s, p>0.9; 40 min: 0.4 ± 0.7 s, p=0.03, 60 min: 0.07 ± 0.03 s, p<0.0001, (Fig. 5 B-C). These data suggest that both K_ir_ channels and GJs coordinate to maintain low potassium levels near nerve terminals and synapses to sustain neuronal excitability during long bouts of activity.

**Figure 5.**
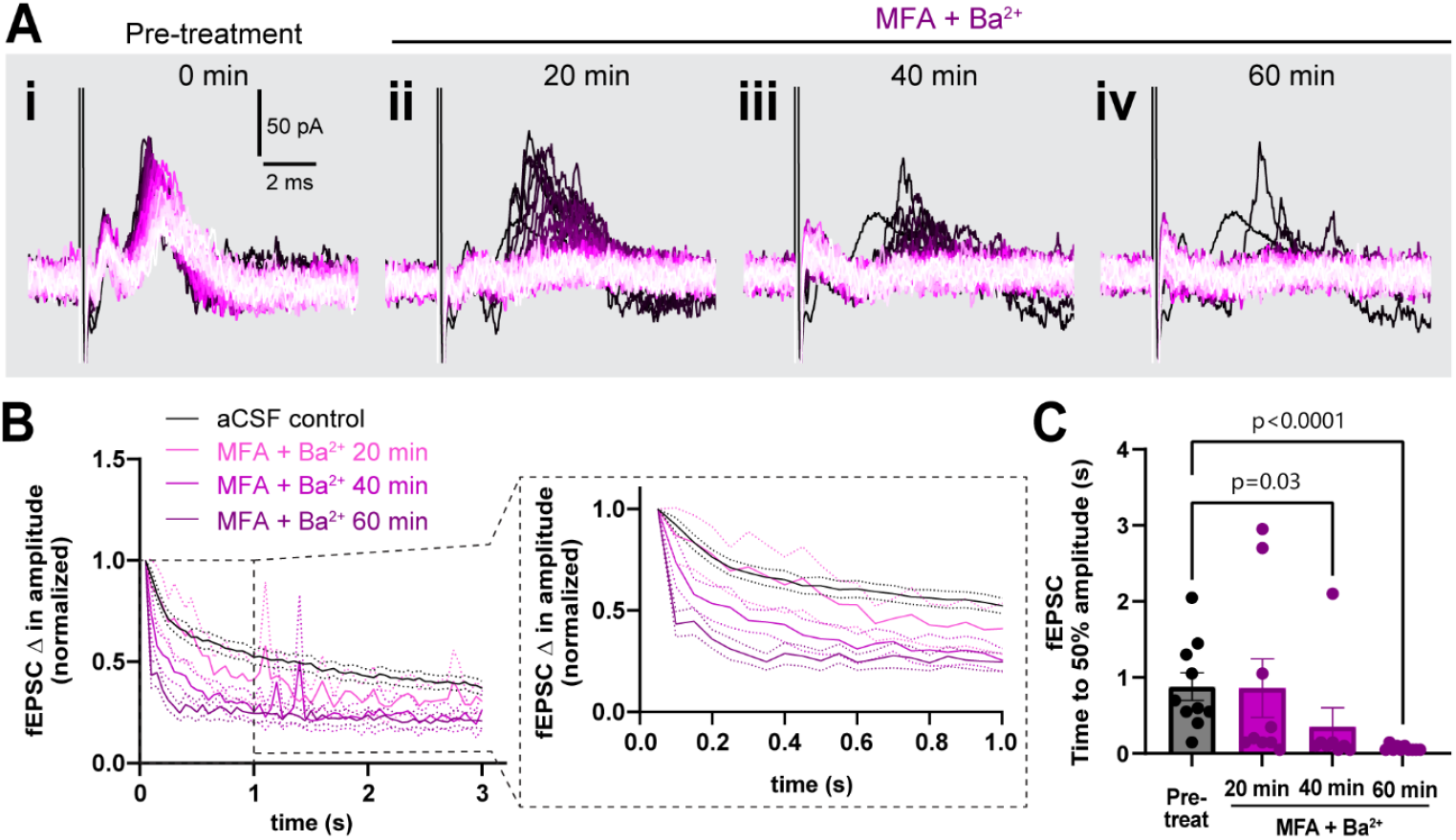
Simultaneous block of GJs and Kir channels progressively worsened sustained neuronal activity. **A**. Representative traces of response progression over 1 hour in MFA + Ba^2+^-treated slices, shown at pre-treatment (i), 20 min (ii), 40 min (iii) and 60 min (iv). **B**. Average amplitude decrease of fEPSCs during 20 Hz stimulation in the pre-treatment (black, n=10), and in MFA + Ba^2+^ after 20 min (pink, n=9), 40 min (light purple,n=8) and 1 hour (dark purple; n=10). Inset on the right shows an expanded view of the amplitude decrease during the first second. **C**. Time to 50% fEPSC amplitude was decreased 40 min and 60 min after co-applying MFA + Ba^2+^ (Kruskal-Wallis test). Data shown as mean ± sem.

## DISCUSSION

Our goal was to evaluate the effect of K_ir_ channels and GJs on neuronal excitability. We measured the effect of acute block of K_ir_ and GJs (with Ba^2+^ and MFA, respectively) on cortical field activity. Our data indicate that both K_ir_ channels and GJs modulate neuronal excitability, presumably by contributing to K^+^ buffering, and that their effect is bidirectionally sensitive to the type of neuronal activity under scrutiny. K_ir_ and GJs can induce hyperexcitability of single evoked responses or sparse activity, as demonstrated by the increased amplitude of the first response, while suppressing sustained responses during prolonged activity, likely due to depolarization of the neuronal membrane^13,14^. Indeed, increases in [K^+^]_e_ produce an initial facilitation of postsynaptic currents due to increased pre-synaptic Ca^2+^ influx and higher neurotransmitter release, but result in inactivation during prolonged stimulation^15^, likely due to neurotransmitter depletion as well as action potential failures and synaptic depression^5^ caused by inactivation of voltage-gated Na^+^ channels. Presumably, applying the blockers individually lead to a moderate [K^+^]_e_ increase, showing an initially hyperexcitable response that could not be sustained during the 20 Hz stimulation. On the other hand, simultaneous treatment with both drugs presumably lead to a higher [K^+^]_e_ increase and much stronger suppression of activity.

K^+^ homeostasis can also be regulated via other pathways not involving K_ir_ and GJs (e.g. Na/K-ATPase pumps on neurons and astrocytes). Given the importance of K^+^ homeostasis for brain health, it is conceivable that functional redundancy exists. The contribution of Na^+^/K^+^ ATPases to K^+^ homeostasis^13,16,17^ and its relationship to K^+^ buffering by astrocytes, however, is still unclear. We observed stronger evoked activity in the first recording of each stimulation train, after an extended period of rest. This hyperexcitability was present in the first few responses even after short periods of recovery during which activity was not exogenously evoked (i.e., the 2-minute intervals between 3 trials for each condition). This suggests that other mechanisms, such as Na^+^/K^+^ ATPases or perhaps K^+^ diffusion, may be sufficient to maintain [K^+^]_e_ within a physiological range during sparse activity, given enough time. However, under sustained activity, as is the case during 3s, 20 Hz stimulation, these systems appear unable to keep up. Notably, cortical neurons fire between 20 Hz and 100 Hz frequencies, underscoring the relevance of the disruption we describe here^18^. The particularly strong effect of the co-treatment with Ba^2+^ and MFA indicates that K_ir_ and GJs together play a major role in maintaining activity of cortical neurons.

### Comparison to prior results

Inhibiting both K_ir_ and GJs together by co-application of Ba^2+^ and MFA strongly disrupted neuronal responses to 20 Hz stimulation. Importantly, we noticed that, when applied individually, each inhibitor took a long time (1 hour) to exert robust disruption of neuronal activity, whereas applying both simultaneously had a robust and rapid effect within only 20-30 min and progressively reduced the response over 1 hour. This observation was surprising, as we expected that once K_ir_ channels are blocked a further block of GJs would not produce an additional affect via the same pathway in buffering [K^+^]_e_. It may indicate that the dose of the inhibitors used (barium and MFA) did not block all the K_ir_ and GJs, as both K_ir_ and GJs are highly abundant on astrocytes^1,19-21^. Nonetheless, it has been reported that 100 μM MFA applied for 1 hour blocked 99.3% of gap junctions^22^, whereas 100 μM Ba^2+^ blocked 48% of K_ir_ conductance in 10 min on patched astrocytes^23^. Co-application potentially increased the efficiency with which the K^+^ buffering system is blocked and thus produced a faster and stronger disruption of neuronal excitability.

The use of genetic knockouts of K_ir_ and GJs has been employed to examine their roles further and this should be attempted in future studies. However, genetic knockout models also are complicated by compensatory or adaptive responses to neuronal firing. This issue seems to be the case for astrocyte-specific K_ir_4.1 cKOs^1^ (using GFAP:Cre), which do not survive past 30 days but exhibit intact action potential firing at P15-P20^1^. In contrast, mice with global Cx30 knockout and astrocyte-specific Cx43 knockout^3^ (also using GFAP:Cre) were reported to look healthy, yet show spontaneous seizures and evoked-hyperexcitability^3^. An inducible conditional astrocyte-specific knockdown of Cx30 and Cx43 (controlled by GLAST:CreERT2) showed that post-synaptic excitability was enhanced despite a decrease in pre-synaptic excitability^2^, similar to our own observations. Yet, this study also showed that deletion of GJs from astrocytes induces widespread gliosis and transcriptomic changes^2^, making it difficult to attribute the changes in neuronal excitability to GJs alone. Our findings based on acute inhibition of K_ir_ and GJs circumvent some of these issues but are, in turn, limited by non-cell-specific action of the inhibitors. Developing cell-specific tools and strategies for disrupting K^+^ buffering in a time-controlled manner will be crucial for further elucidating the role of astrocyte GJs and K_ir_4.1 in regulating K^+^ homeostasis.

### K^+^ homeostasis: spatial or electrical buffering?

Separate reports of K_ir_ and GJ contribution to K^+^ buffering may not have accurately captured K_ir_ and GJ interaction, leading to incomplete hypotheses and models of K^+^ buffering^13,24^. The interaction between K_ir_ and GJs largely explains passive electrical properties of mature astrocytes and is necessary for K^+^ buffering, as shown by experimental data and computational modeling^23,25^. Astrocytes are an ideal candidate for buffering K^+^ because of two key features set by K_ir_ channels and GJs: 1) K_ir_ (alongside other K^+^ channels) sets the resting membrane potential of astrocytes, hyperpolarizing them by bringing the membrane potential values close to E_k_. As a result, during neuronal firing, astrocytes are highly sensitive to [K^+^]_e_ increases, which is why they depolarize (albeit to a much smaller degree) in synchrony with neurons^26^; 2) coupling of astrocytes via GJs provides an “electrical buffering” system and equilibrates the membrane potential^22^, keeping depolarizations to a minimum in the face of large increases of K^+^ uptake. The role of GJs in allowing currents to leak between coupled cells, thus stabilizing resting membrane potential and dampening excitability, is a property that is well documented for electrical synapses among interneurons^27^. Importantly, in astrocytes, GJs help maintain their relatively hyperpolarized membrane potential, which sustains the driving force for continuous K^+^ removal from the extracellular space. Without GJs, K_ir_-mediated K^+^ entry would produce large depolarizations of astrocytes and impair their capacity to take up K^+^ during neuronal firing^25,28^. This framework suggests that GJs contribute to K^+^ buffering not as much by physically moving K^+^ between astrocytes (spatial buffering) but rather by maintaining their membrane potential hyperpolarized to sustain the driving force for K^+^ uptake (electrical buffering). Thus, perhaps our theoretical framework should be revised to think of the astrocyte syncytium less as a spatial buffering system and more as an electrical buffering system.

Our observation that K_ir_ and GJs contribute similarly to neuronal excitability and strongly impair neuronal activity when blocked together aligns with this framework; larger astrocyte depolarizations would lead to stronger disruption of [K^+^]_e_ homeostasis and, therefore, of neuronal activity. Existing discussions of astrocyte-mediated K^+^ buffering often do not appropriately account for GJs and thus assume that K^+^ entry-mediated depolarizations are bigger^13^. Consequently, they underestimate the buffering capacity of astrocytes. In contrast, accounting for K_ir_ and GJs in the way we propose is in line with the idea of the development of electrical buffering system and the electrically passive feature of the astrocyte syncytium^22,23^. Indeed, these passive properties may be exactly what allows astrocytes to have an active and continuous role in regulating neuronal excitability.

## Conclusion

In summary, we show that K_ir_ channels and GJs bidirectionally regulate neuronal activity, whereby they restrain hyperexcitability during sparse firing conditions but prevent neuronal fatigue during sustained activity. We propose that this is largely due to the passive properties endowed on astrocytes by GJs, thus preserving a constant drive for K^+^ influx through K_ir_ channels and hence maintaining [K^+^]_e_ within physiological bounds to sustain neuronal activity. Understanding how disruptions of passive properties due to changes in K_ir_ channels or GJs on astrocytes affect neuronal circuits could provide valuable insights into brain disorders with disrupted neuronal excitability, such as epilepsy^29^, ischemia^30^, and autism^31^.

## ACKNOWLEDGMENTS

The work was sponsored by the American Heart Association Predoctoral fellowship 23PRE1022986 (DB), and National Institutes of Health R01 NS110690 (AM) and R01 DC012938 (HvG). We thank Bradley Marxmiller for his invaluable help in troubleshooting experiments and equipment. We also thank Dr. Gary Westbrook for his advice on experimental setup and analysis, and for reading the manuscript.

## AUTHOR CONTRIBUTIONS

DB, HvG and AM conceived of the project and designed experiments. DB conducted all experiments and analyzed data. AD and SJS helped develop analysis pipelines. DB drafted the manuscript and DB and AM prepared the figures, with edits from all co-authors. DB, HvG and AM finalized the manuscript.

