## Supplemental Figures for "Astrocyte gap junctions and Kir channels contribute to K^+^ buffering and regulate neuronal excitability"

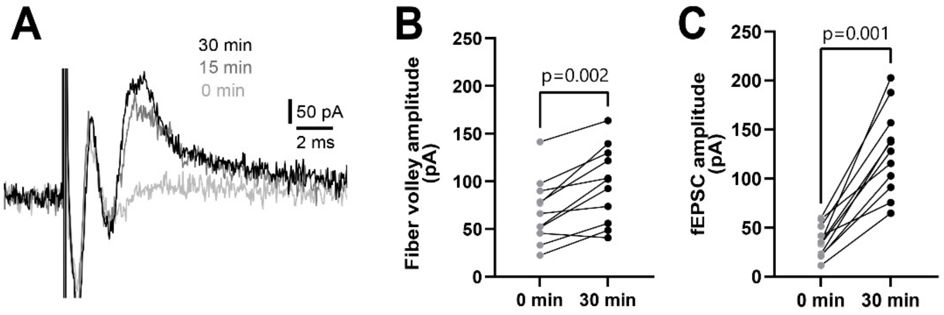


**Supplemental figure 1.** Response stabilization over 30 min. **A.** Representative traces showing the increase in field response amplitude over 30 min. **B.** Both fiber volley (left) and fEPSC amplitude (right) increased over 30 min of stabilization (n=10, Wilcoxon test).

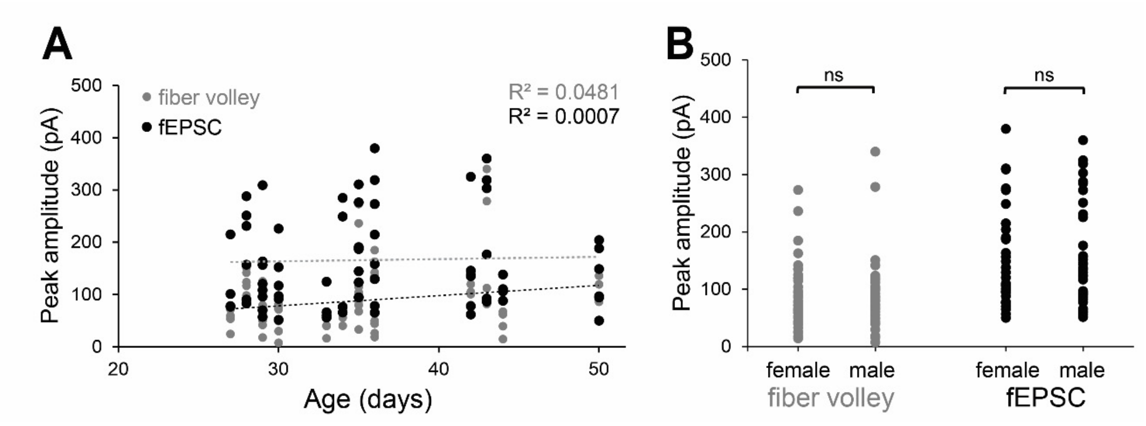


**Supplemental Figure 2.** Peak amplitude variability is not explained by age or sex. **A.** There was no correlation between mouse age and peak amplitude of either fiber volley or fEPSC (n=37, linear correlation). **B.** There was no difference in peak amplitude of either fiber volley or fEPSC between males and females (n=8 female, 18 slices; n=9 male, 19 slices; Mann-Whitney test).

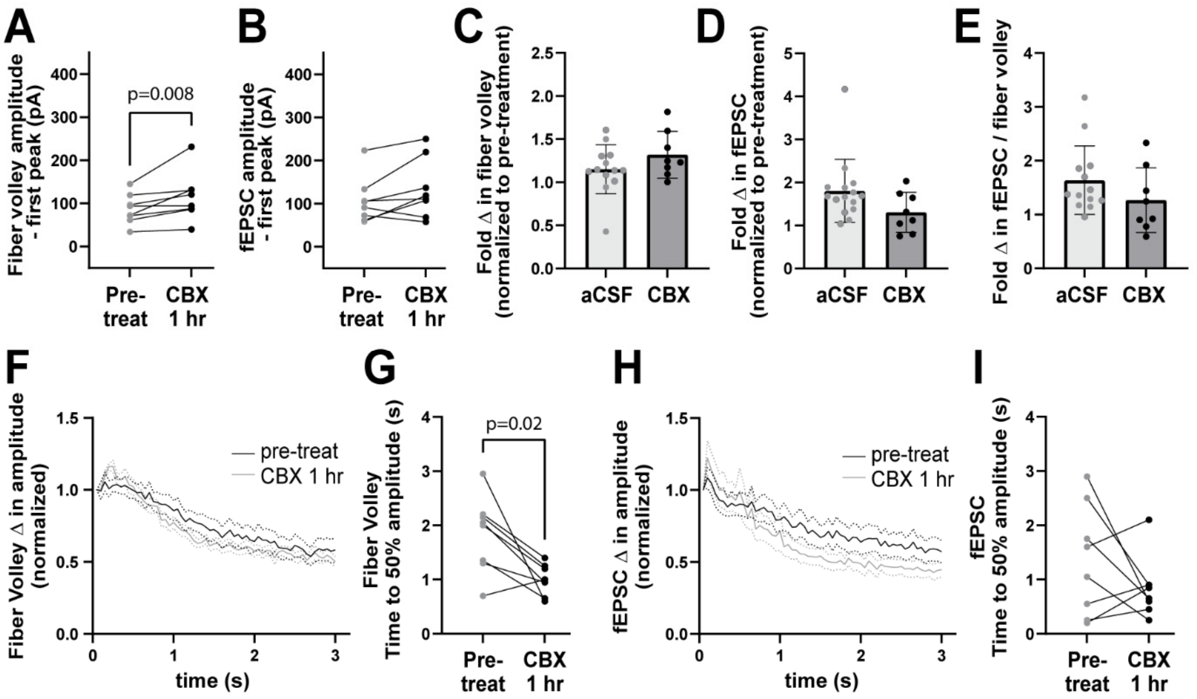


**Supplemental Figure 3.** Carbenoxolone (CBX) had a mild effect on neuronal responses. **A-B.** Fiber volley amplitude (**A**) was increased after 1 hour in CBX while fEPSC amplitude was unchanged (n=8, Wilcoxon test). **C-E.** The fold change in the amplitude of fiber volley (**C**) and fEPSC (**D**), compared to pre-treatment values, as well as the fiber volley/fEPSC ratio (**E**) were unchanged in CBX (n = 8) compared to aCSF control (n = 15) after 1 hour (Mann Whitney test). **F-G.** Amplitude decrease of fiber volley over the 20 Hz stimulation (**F**) showed a slightly faster time to 50% (**G**) after 1 hour in CBX (n=8; Wilcoxon test). **H-I.** Amplitude decrease of the fEPSC over 20 Hz stimulation (**H**) and time to 50% (**I**) showed no difference after 1 hour in CBX (n=8; Wilcoxon test). Data shown as mean ± sem.
